# Ancestral Reconstruction of Protein Interaction Networks

**DOI:** 10.1101/408773

**Authors:** Benjamin J. Liebeskind, Richard W. Aldrich, Edward M. Marcotte

## Abstract

The molecular and cellular basis of novelty is a major open question in evolutionary biology. Until very recently, the vast majority of cellular phenomena were so difficult to sample that cross-species studies of biochemistry were rare and comparative analysis at the level of biochemical systems was almost impossible. Recent advances in systems biology are changing what is possible, however, and comparative phylogenetic methods that can handle this new data are wanted. Here, we introduce the term “phylogenetic latent variable models” (PLVMs, pronounced “plums”) for a class of models that has recently been used to infer the evolution of cellular states from systems-level molecular data, and develop a new parameterization and fitting strategy that is useful for comparative inference of biochemical networks. We deploy this new framework to infer the ancestral states and evolutionary dynamics of protein-interaction networks by analyzing >16,000 predominantly metazoan co-fractionation and affinity-purification mass spectrometry experiments. Based on these data, we estimate ancestral interactions across unikonts, broadly recovering protein complexes involved in translation, transcription, proteostasis, transport, and membrane trafficking. Using these results, we predict an ancient core of the Commander complex made up of CCDC22, CCDC93, C16orf62, and DSCR3, with more recent additions of COMMD-containing proteins in tetrapods. We also use simulations to develop model fitting strategies and discuss future model developments.

## 1. Introduction

Evolutionary biology has largely concerned itself with the analysis of phenotypic traits and molecular sequence data. These sorts of data are relatively easy to collect as compared to the cellular and molecular traits that are causally intermediate between genotype and phenotypes. The difficulty of collecting these intermediate data, on the other hand, has historically limited the taxonomic reach of many areas of biology. Recently, however, a variety of new methods are breaking down this old divide and opening up the cell to large-scale, systematic data collection. Crucially, many of these new techniques are not reliant on traditional model organisms. These methods can probe the state of the cell across its hierarchical organization, including: functional states of the genome, such as transcription factor binding (ChIPseq) [1], genomic architecture (Hi-C) [2], epigenetic modifications [3], and replication timing (repli-Seq) [4]; events in the transcription/translation process, including single-cell quantitative gene expression [5], splicing (spliceosome capture) [6], and translation (ribosome capture) [7]; and increasingly comprehensive protein interaction maps that capture functioning proteins in their native environment [8, 9, 10].

These new data types are a boon to evolutionary biologists. Ever since Darwin “contemplate[d] a tangled bank, clothed with…elaborately constructed forms, so different from each other, and dependent upon each other in so complex a manner,”[11] it has been recognized that evolution is directed both by invariant fundamental laws and by the ever-changing environment in which those laws play out. The evolution of a protein within the cell and the evolution of species within its environment are the same in this regard: both are contingent on the structure of their immediate context. Evolutionary biologists are therefore interested in the evolutionary dynamics not just of the replicating units of evolution - species, phenotypes, genes - but also of the relationships between these units; their interaction networks [12, 13, 14]. Indeed, many questions of immediate interest to evolutionary biologists, such as the origins of complex tissue types, are unanswerable without detailed knowledge of the evolution of molecular systems [15]. With the advent of so many new data collection methods, comparative analysis of the molecular ecology of the cell is now a real possibility. Indeed, many have already been used to elucidate evolutionary processes [16, 17, 18, 19].

We have focused our efforts on protein interaction networks. Most proteins function in groups, and many of the core functionalities of cells, such as transcription, translation, and splicing, are carried out by large, stable, multiprotein complexes. Proteomics experiments that report on global protein interaction networks are largely of two sorts: affinity-purification mass spectrometry (AP-MS), where bait proteins are isolated with antibodies, taking presumed interactors (“prey”) with them; and co-fractionation mass spectrometry (CF-MS), where native biochemical fractionation is used to separate cell lysate and the co-fractionation of protein pairs is used as a read-out of interaction strength. Whereas large-scale AP-MS is only possible in species amenable to genetic tagging, CF-MS is broadly applicable across species, with one recent study collecting data from nine eukaryotes [9]. These proteomics datasets can be used to answer several longstanding questions about molecular evolution, for instance: What are the evolutionary dynamics of protein-interaction networks? What kinds of architectural changes in the network are responsible for the emergence of novel cellular phenotypes? How does evolution mitigate the risk of protein-network dysfunction arising from deleterious mutations?

To leverage these new data types, it is necessary to have suitable models. Supervised machine learning has become the standard modeling strategy for protein interaction data [8, 20, 21], and recent studies have extended this approach to cross-species data by combining experiments from several species in a single feature matrix on the basis of orthology [9, 8]. Machine learning excels at extracting patterns from noisy, high-dimensional data, and this approach successfully leverages evolutionary conservation to discovery of conserved protein interactions, but because it does not use phylogenetic information it has two disadvantages from the perspective of evolutionary biologists. First, by ignoring phylogenetic relatedness when adding species to the feature matrix, it runs into statistical problems first identified by Felsenstein [22], though we note that the machine learning techniques commonly applied are fairly robust to non-independent features. But more importantly, this technique cannot recover evolutionary history. To do so, evolutionary biologists commonly use stochastic Markov models of evolution that can probabilistically resolve interior states of the phylogeny. However, these have the disadvantage of requiring reliable data measurements at the tips of the phylogeny, and unlike nucleotide sequencing, the inference of interaction networks entails substantial uncertainty. The first methods used to infer the evolution of protein interaction networks outsourced this difficulty by taking preprocessed networks as input [23], but more sophisticated methods have been developed that model measurement uncertainty directly.

The most promising of these methods models changes in the network as a discrete-state Markov model, just as in standard phylogenetic comparative methods. This discrete component is then coupled to a model that describes the expected distribution of observable data given the unobserved state at each leaf of the tree. Several different formulations have been suggested, almost all of which are designed for gene expression data [24, 25, 26]. We introduce a general term for these types of models: phylogenetic latent variable models (PLVMs, pronounced “plums”). (Note that PLVMs are distinct from “tree-HMM” models, where the latent variable is a linear Markov process, with each state being associated with a set of tree parameters [27, 19]).

Existing implementations of PLVMs are promising, but not easily applicable to protein-interaction data. The most sophisticated PLVM implementations are Arboretum [24] and MRTLE [26], two related methods from the Roy and Regev labs. These models detect gene co-expression modules from expression profiles, which must have the same dimensionality across species, by modeling module membership as the latent state. While powerful for gene expression data, our protein interaction datasets do not have identical dimensionality across species, and for network inference it is more desirable to model the network edges as the latent state. Two other implementations, ProPhyC [28] and tHMM [29], also have disadvantages. In ProPhyC, the error model requires a fixed parameter for every edge in the network, ancestral reconstruction is not probabilistic, and it can only consider a single data source. Unlike ProPhyC, tHMM does use probabilistic ancestral state reconstruction, but is only implemented for one character at a time and for one data source. Finally, all current studies on PLVMs fix certain parameters in the model before fitting, and Arboretum, MRTLE, and ProPhyC use an expectation-maximization algorithm, a local fitting procedure. This raises the question of whether it is possible to obtain good fits for PLVMs on the input data, or whether the user must use extraneous data to estimate parameters, as in current implementations.

We therefore set out to develop a broadly applicable implementation of a PLVM, with the following features: 1.) Probabilistic ancestral state reconstruction of networks 2.) Global fitting procedure 3.) Ability to leverage different data sources 4.) Capable of handling genome-scale data 5.) Flexible and modular model specification. Below we describe such an implementation, its performance on simulated and real protein-interaction datasets, and the resulting reconstruction of ancestral protein interaction networks.

## 2. Results

### 2.1. A Generalizable Phylogenetic Latent Variable Model

Our model takes as input a time-calibrated phylogeny and a data matrix comprising one or more features that report on the presence of an interaction for every pair of proteins under consideration, and returns probabilities of an interaction at every node of the input phylogeny. It contains two parts: a discrete state continuous time Markov chain (CTMC) along the phylogeny modeling the gains and losses of interactions, and an error model mapping these latent states to observed data. The CTMC component is parameterized by two instantaneous rate parameters, *α* and *β*, that describe the rate of gains and losses, respectively. The error model is two multivariate Gaussian distributions, *N* (*µ*_0_, Σ_0_) and *N* (*µ*_1_, Σ_1_), that describe the expected distributions of the data features given the non-interacting (0) and interacting (1) states (Figure 1). The likelihood of each state under the two error models given the input data begins the calculation of a tree likelihood from the CTMC using Felsenstein’s pruning algorithm [30]. Ancestral states are inferred in a similar fashion, with the addition of a root-to-tip traversal (See extended methods).

**Figure 1:**
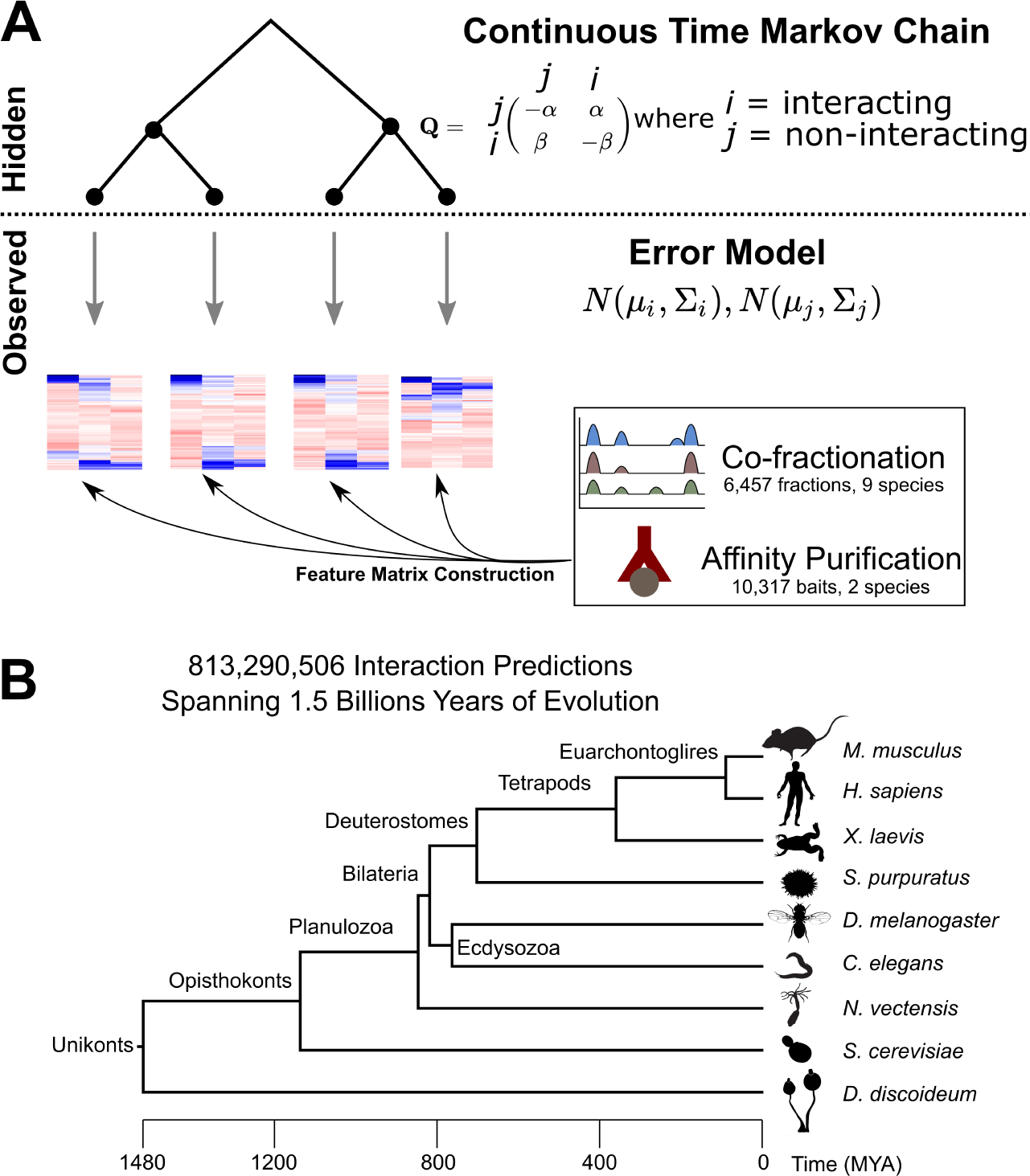
Modelling Framework. **A.)** Schematic of the phylogenetic latent variable model, with a continuous time Markov chain describing the evolution of network edges and an error model mapping the unobserved edge states to continuous, observed data. Input features were extracted from several high-throughput datasets, comprising over 16k mass spectrometry experiments **B.)** Time-calibrated phylogeny used for inference.

This framework is simple, flexible, and powerful. It uses few parameters relative to other extant PLVMs, but can handle features coming from very different data sources – in our case, both AP-MS and CF-MS proteomics experiments. It deals with missing data in a principled way, and is also extensible to large datasets. While we use the model to predict protein-interaction networks, it is applicable in principle to any kind of network data. We use simulated annealing, a heuristic search algorithm, to fit the model on training subsets of the data before predicting on the full dataset. Our final predictions cover >800 million network edges spanning the 17-node tree of animals and two eukaryote outgroups (Figure 1B). The model and implementation is available in our Python package (https://github.com/marcottelab/plum). This package supports several other error models besides the multivariate Gaussian described here, including Gumbel and Cauchy distributions for univariate data and diagonal multivariate Gaussians, and is designed to make it easy for developers to add their own models.

### 2.2. Simulations and model performance

Published studies on PLVMs fix either the error model parameters, the CTMC parameters, or both, before fitting the model and using it for prediction [25, 24, 26]. Is it possible to fit the entire model and achieve good performance? Protein interaction networks are a good testbed for these models because large curated datasets of protein-protein interactions exist in a number of species [31, 32]. Interactomic studies using other modeling strategies have found that, while current methods are a vast improvement on older methods, such as yeast two-hybrid, they are still very noisy [8]. Specifically, weak signal, high false-negative rates, and class imbalance (non-interacting pairs far outnumber interacting pairs), all plague statistical predictions. For instance, the distributions of one feature, Pearson’s correlation coefficient calculated on fractionation profiles of gold-standard human protein-interactions, is barely distinguishable from the distribution among negative interactions (Figure 2A).

**Figure 2:**
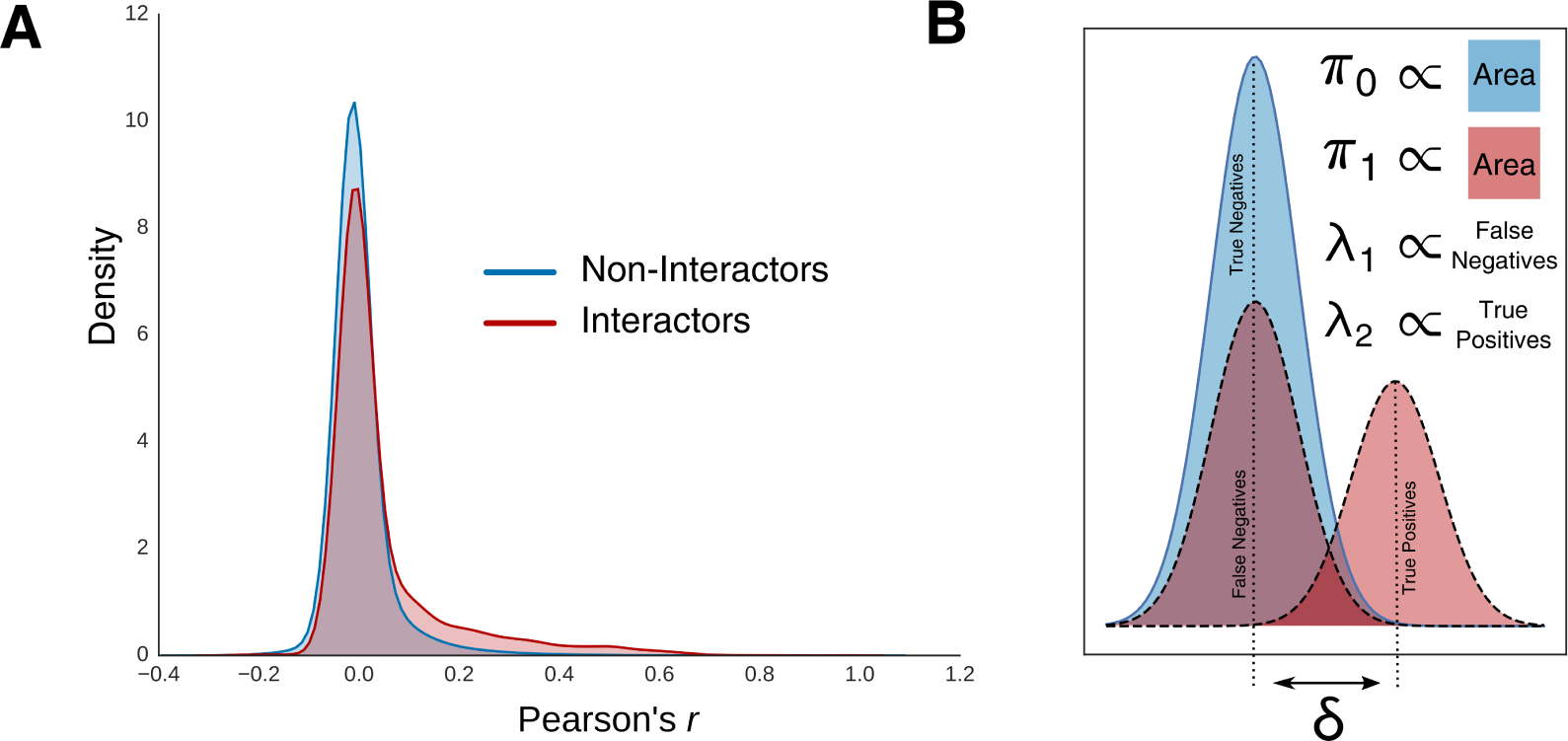
Simulation Strategy. **A.)** The distribution of Pearson’s correlation coefficient averaged over 32 co-fractionation experiments on human cell lines. The distribution for gold-standard interacting human protein pairs is barely distinguishable from non-interactors. **B.)** Error models parameters were manipulated to replicate typical experimental noise. The equilibrium frequencies of the two state, *π*0 and *π*1, can be tuned to replicate class-imbalance. The difference between the means *δ*, was changed to replicate weak signal. The mixture weights for the positive error model, *λ 1* and *λ 2*, set the false negative rate.

To assay model performance in a controlled fashion, we simulated data under the PLVM in a way that replicates the common biases and difficulties of protein interaction data (Figure 2B), and tested different fitting strategies. To replicate false negatives, we used a different error model than that used for prediction. Instead of a single multivariate Gaussian for the interacting state, we simulated under a two-component mixture given by 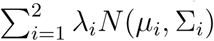, and fixed the mean vector of the first component to equal that of the Gaussian assigned to the negative state. This allowed us to change the false negative rates by tuning the mixture weights (*λ*_*i*_). As *λ*_1_ approaches 1, the distributions of positive- and negative-labeled data becomes indistinguishable. We also simulated over different values of *α* and *β*. These parameters by themselves determine the rate of evolution, and the equilibrium frequencies of the two states, *π*_0_ and *π*_1_, which are the normalized rate parameters, 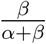and 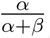, respectively, determine the class imbalance because they are the expected frequencies of the two states after an infinitely long Markov chain.

Finally, we simulated over different distances between the means of the positive and negatively labeled data. In all, we simulated datasets from 360 different parameters combinations, with 10 replicates each.

The phylogeny used for simulation and inference was obtained from Time Tree (http://www.timetree.org/) [33].

We fit the model to datasets from each parameter combination and predicted the states at the tips of the tree on a different replicate as well as on the training data. Model performance was evaluated by comparing the trade-off between precision and recall using the average precision score (APS). In general, fitting the model by maximum likelihood performed poorly by this measure on both training and test data (Figure 3A). We then tried fitting the model using the APS itself as a criterion. Unsurprisingly, this strategy far outperformed likelihood on training data, but more importantly, it performed better on hold-out data as well, suggesting that PLVMs may be more useful when implemented in a supervised learning framework using empirical training data (Figure 3A).

**Figure 3:**
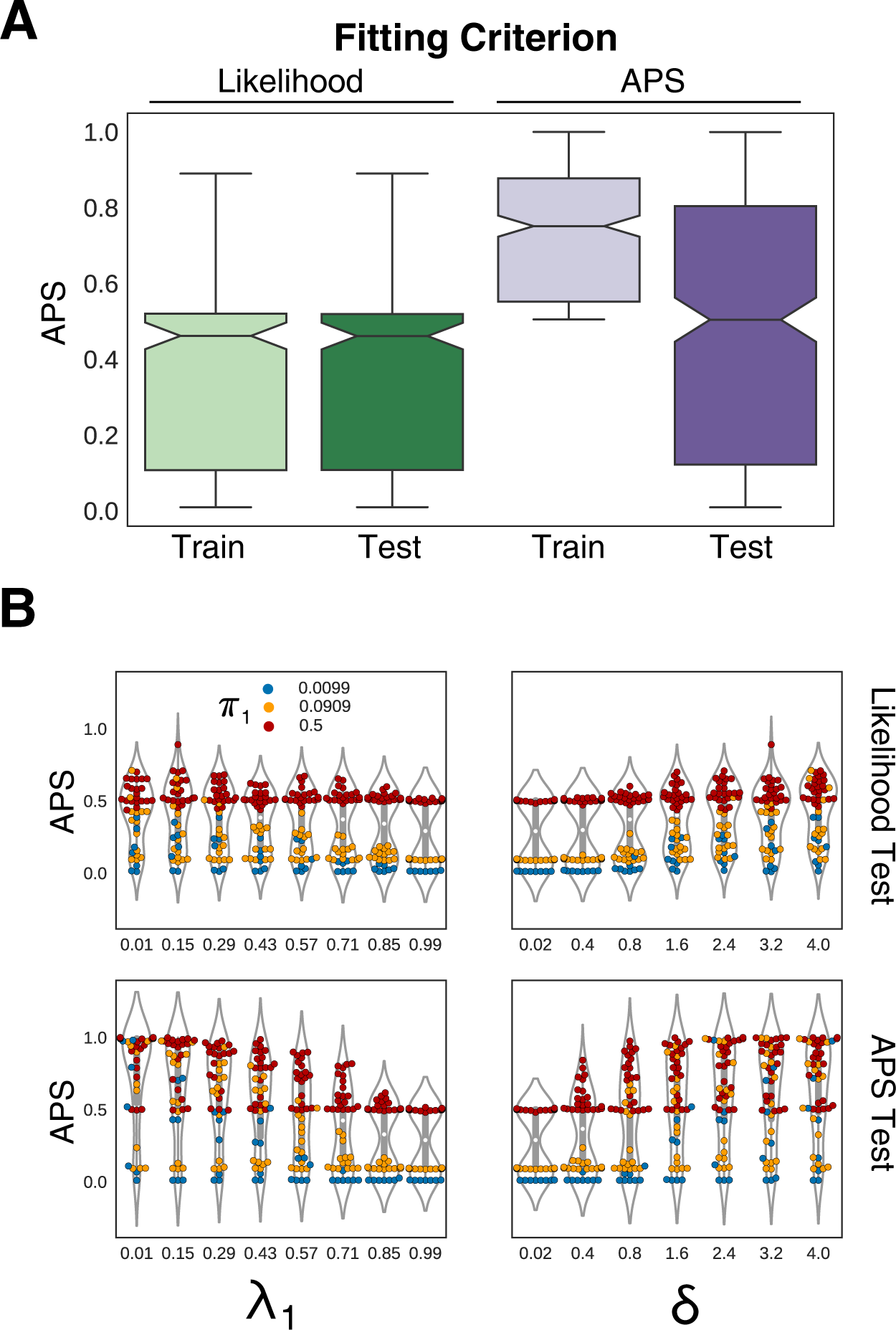
Simulation Results. **A.)** Performance on simulated training and test (hold-out) data when the model is trained by maximizing either the likelihood or the average precision score (APS). Fitting by APS outperforms fitting under likelihood. The APS is also used as a criterion for goodness of fit and results include all simulation parameter combinations **B.)** Performance as a function of the mixture weight *λ 1*, the false negative rate, *δ*, the distance between the means of positive and negative interactions, and the equilibrium frequency *π* 0 which is proportional to class imbalance. *π* 0has a surprisingly large effect on performance.

What parameter most affected model performance? Unsurprisingly, increasing the false negative rate by increasing the mixture component *λ*_1_ hurt performance in both fitting strategies, as did weakening the signal by decreasing the distance *δ* between the means of the positive and negative error models. Perhaps more surprisingly, the largest single factor seems to be class imbalance, as measured by the equilibrium frequencies. When *λ*_1_ and *δ* are in unfavorable regions of parameter space, the performance of the model is determined entirely by the class imbalance, and even in the best regions of the other parameters, a strong class imbalance can significantly hurt performance (Figure 3B).

### 2.3. Performance on hold-out sets

The availability of curated protein-interaction data sets from several of our included species provide an opportunity to test modeling strategies on real data that was withheld from training. We found that the model is able to recapitulate known protein interactions across species even when relatively little data is available for that species, as in mouse, which is represented by only two fractionation experiments (Table 1) and was not used for training (Figure 4A). To quantify the effect of the model, we plot the performance of the raw features collected directly from the data in each species individually alongside the model precision-recall curves. As expected due to its low coverage, the model dramatically improves performance in mouse, but it also does so in humans, which has the most data for any lineage, showing the power of comparative methods. Fly and yeast are separated from other species by much deeper branches than human or mouse, and correspondingly are improved less by the model. Interestingly, though the large AP-MS dataset in yeast [34] performs strongly on its own, the addition of the model improves performance in the high-precision/low-recall regime where the AP-MS data does poorly, but at the cost of overall recall.

**Figure 4:**
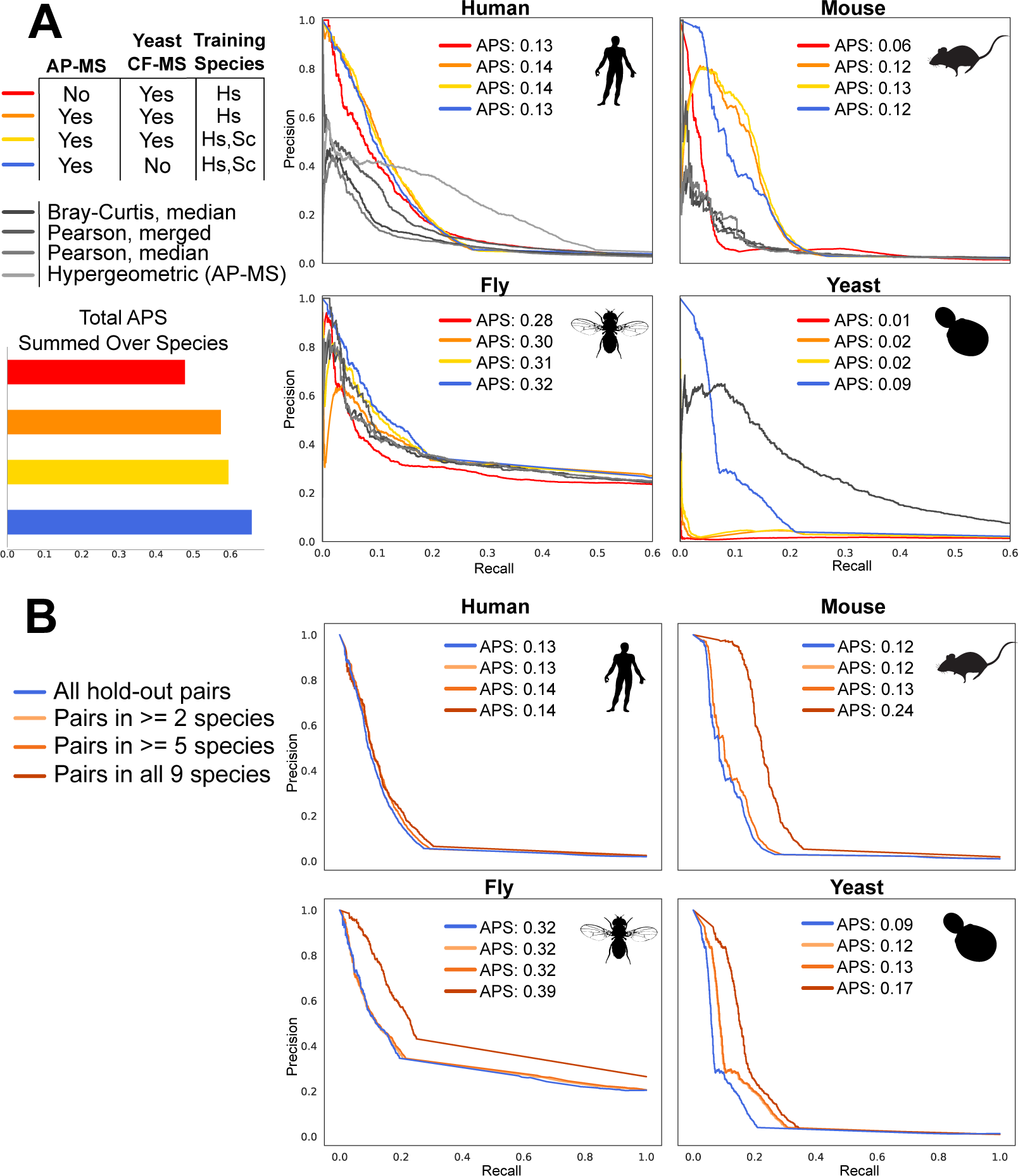
**A** Performance on hold-out sets in four species, measured as precision-recall curves and the average precision score (APS). Three modeling conditions are plotted next to the raw features derived individually in each species from the highest performing (blue) dataset. This dataset was also used for all subsequent analyses. Note that not all features were collected for each species. The higher baseline in flies is due to a lower ratio of negatives to positives in the test data (see methods), not better performance in that species, and in general the species cannot be directly compared to each other due to differences in the test sets. **B** Conserved orthogroup interactions, where the orthogroups are shared across more taxa, perform better.

As expected, the performance of the model is affected by the nature of the included data, especially in species with less available data (Figure 4A). Adding AP-MS data, for instance, generally improves performance. Unexpectedly, however, we found that *removing* co-fractionation data from yeast and using only AP-MS data substantially improved performance for that species, which was otherwise extremely poor, even when including yeast samples in the training data (Figure 4A). This is likely due to the fact that yeast is represented by only a single co-fractionation experiment, the worst sampling for any species. A single co-fractionation experiment ensures a high false positive rate because there are far fewer fractions than proteins, and because our model requires the same error model for every leaf on the tree, there is no way to down-weight the yeast data relative to other, more fully sampled species. Removing the yeast CF-MS data had a more minor affect on other species and yielded the best overall performance, judged as the average precision score summed over hold-out data for all four species (Figure 4A). This model and its predictions were used for all subsequent observations below.

We also observed that, perhaps unsurprisingly, orthogroups that are shared among more species perform better than those that are more taxonomically restricted (Figure 4B). Orthogroups that are not shared among species cannot benefit from data in those other species and so the main source of the model’s power is absent. However, we note that it is also probably the case that large, stable protein complexes tend to be more highly conserved and so may contribute this result as well.

### 2.4. Reconstructing ancestral networks

Using our best performing model (Figure 4A, blue), we reconstructed ancestral orthgroup interaction networks across the tree. When creating networks from pairwise scores, one typically calculates a false discovery rate (FDR) threshold to trim the network for visualization and further analysis, such as clustering. These steps are not as straightforward in our case because we do not have training data for every node of the tree. How should FDR calculations be propagated across the tree? For a model trained under likelihood, this is straightforward because the model returns probabilities that are directly comparable. But because we train the model using the average precision score as a metric, the scores it returns are not comparable. Indeed, we found that the scores at each node can have quite different distributions.

We therefore converted the scores at each node to z-scores, calculated the false discovery rate using the human hold-out data, and trimmed all the node z-scores at the 25% FDR level. This allowed us to reconstruct sensible networks at each node of the tree. In Figure 5, we show the clustered network of the most recent common ancestor of planulozoans (cnidarians + bilaterians). As expected, we find many of the well known soluble protein complexes that make up the central machinery of mammalian cells were present in this “ur-planulozoan.” Membrane proteins were far less resolved, as is often the case in proteomics due to the difficulty of solubilizing these proteins for experimentation [35], and group together in a large, dense network (Figure 5).

**Figure 5:**
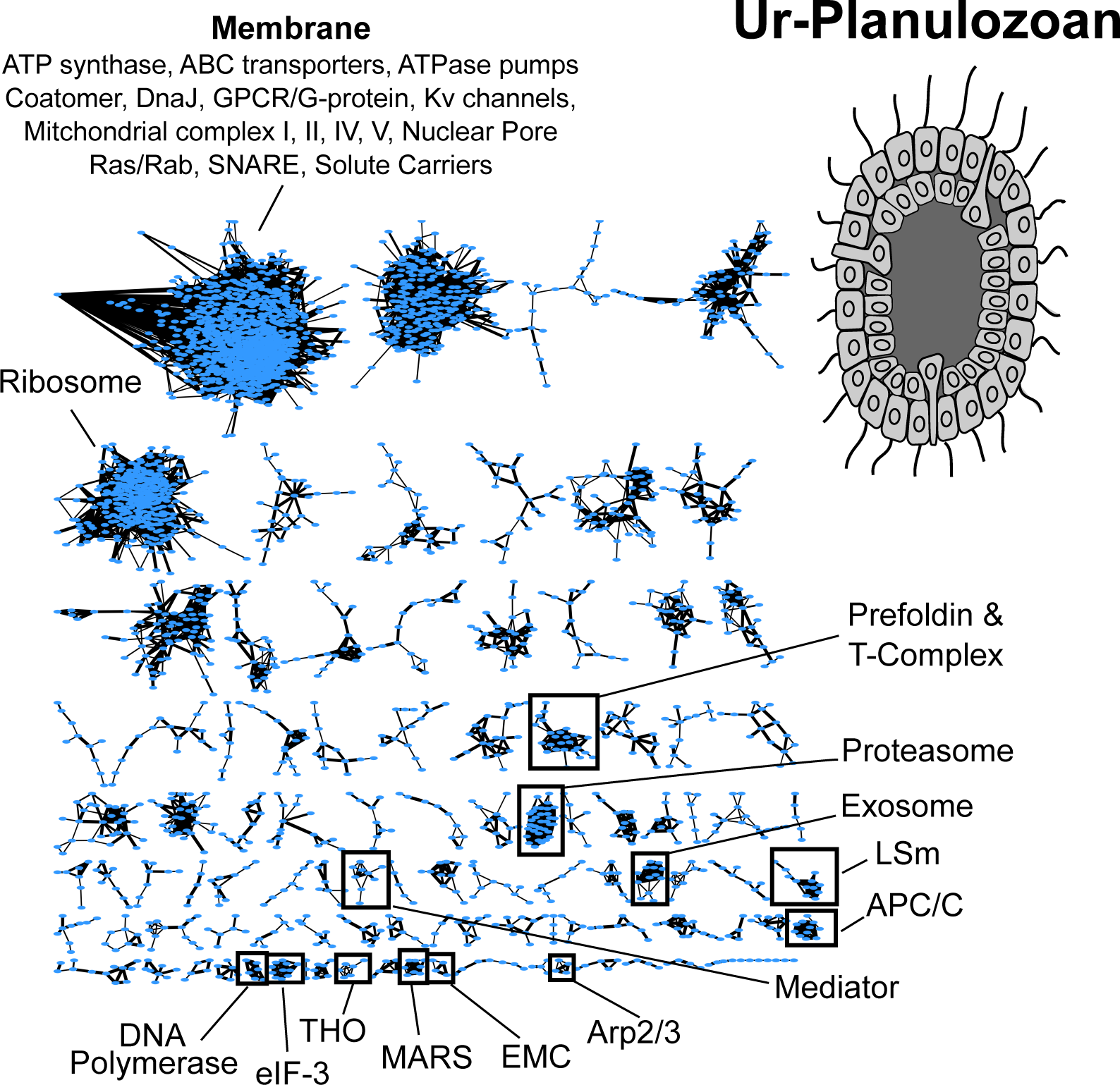
Reconstructed orthogroup interaction network for the most recent common ancestor of planulozoans (cnidarians + bilaterians). The model successfully reconstructs known soluble complexes and groups membrane proteins into a large clump. Edge widths are proportional to the z-score transformed score from the PLVM

**Table 1.**
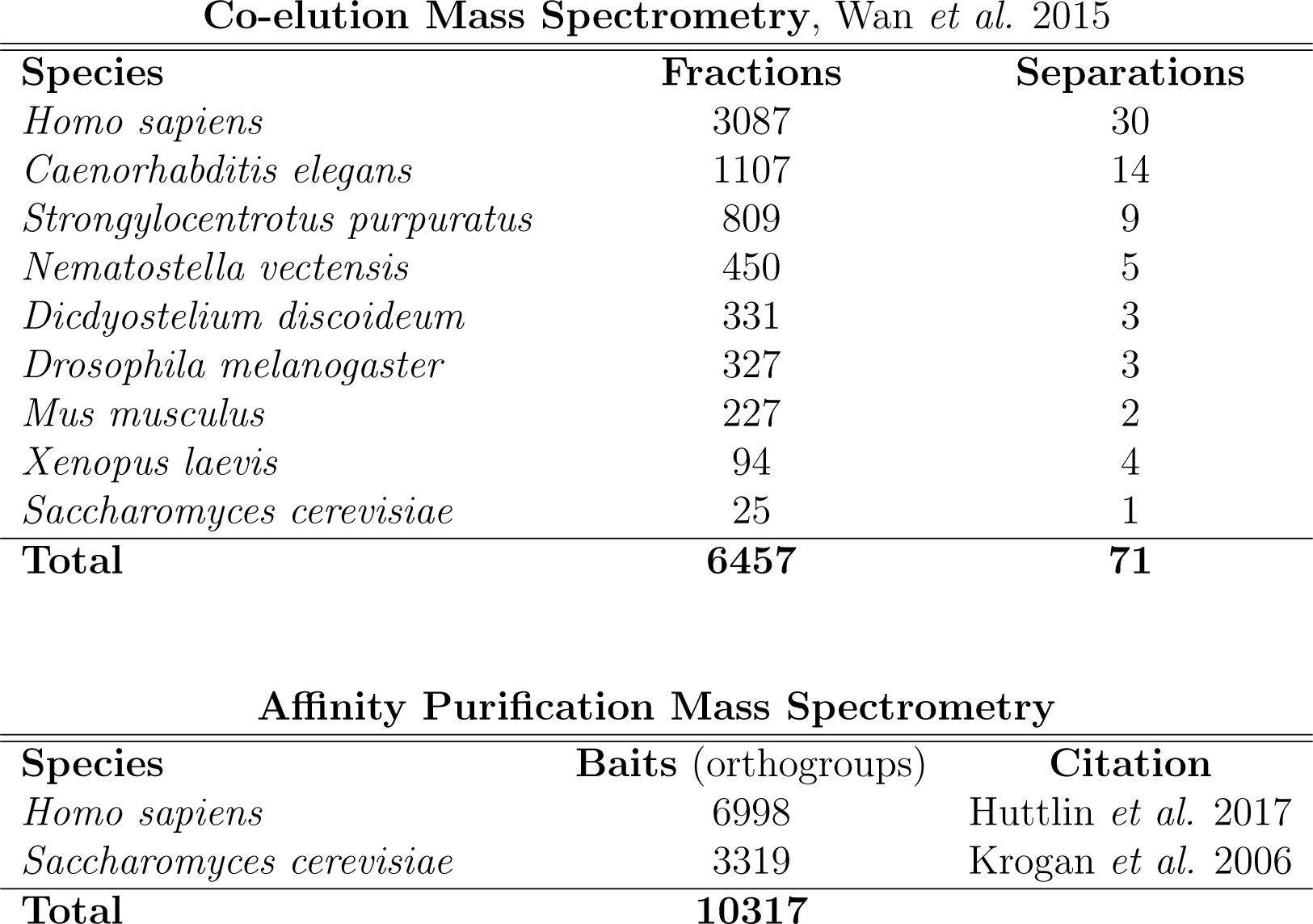
Experimental data used for analyses

Co-elution Mass Spectrometry, Wan *et al.* 2015
| Species | Fractions | Separations |
| --- | --- | --- |
| *Homo sapiens* | 3087 | 30 |
| *Caenorhabditis elegans* | 1107 | 14 |
| *Strongylocentrotus purpuratus* | 809 | 9 |
| *Nematostella vectensis* | 450 | 5 |
| *Dicdyostelium discoideum* | 331 | 3 |
| *Drosophila melanogaster* | 327 | 3 |
| *Mus musculus* | 227 | 2 |
| *Xenopus laevis* | 94 | 4 |
| *Saccharomyces cerevisiae* | 25 | 1 |
| **Total** | **6457** | **71** |

Table 1: Experimental data used for analyses
| Species | Baits (orthogroups) | Citation |
| --- | --- | --- |
| *Homo sapiens* | 6998 | Huttlin *et al.* 2017 |
| *Saccharomyces cerevisiae* | 3319 | Krogan *et al.* 2006 |
| **Total** | **10317** |  |

### 2.5. Evolution of the Commander complex

While it is clear that the PLVM is powerful enough to recover known protein complexes in extant species and reconstruct sensible ancestral networks, this modeling framework is primarily of interest for its ability to reconstruct the evolutionary dynamics of protein complexes. As one example of this application, we explored the evolution of Commander, a protein complex identified in high-throughput proteomics studies of human cell lines and by bioinformatic approaches [36]. Commander was named for the copper metabolism gene MURR1 domain (COMMD) containing family of proteins that make up most of its members. It is involved in endosomal trafficking of a variety of receptors and is associated with several severe developmental disorders. Targeted biochemical studies of Commander subunits, also in human cell lines or mice, have identified several sub-complexes, most notably the CCC complex, composed of COMMD1, CCDC22, CCDC93, and C16orf62, which has been shown to cooperate with the WASH complex in endosomal sorting and targeting of the low-density lipoprotein receptor [37] and the copper transporter ATP7A [38]. Based on this experimental evidence together with structural homology to Retromer, another endosomal trafficking complex, Mallam *et al.* (2017) proposed that Commander recognizes specific cargo proteins and, *via* its interaction with WASH, transduces this information into structural changes in the endosomal vesicle and movement of vesicle along the cytoskeleton. The COMMD-containing proteins themselves may interact with the rest of the complex heterogeneously, creating unique combinations that recognize different cargos.

The proteins constituting the Commander complex go back to the common ancestor of eukaryotes and are broadly retained in vertebrates [36]. Various COMMD-containing proteins have been lost in protostomes, while CCDC93, CCDC22, and C16orf62 are conserved in all animals studied here. Yeast has lost the complex in its entirety, as have most other eukaryotic lineages, though *Dictyostelium dicoideum* has retained it. We therefore wanted to know what evidence there was for the evolution of interactions among these subunits. Did the common ancestor of all Unikonts have a functioning Commander complex?

We found evidence for an ancient interaction between CCDC22, CCDC93, C16orf62, and perhaps DSCR3 (Figure 6). This corresponds most closely to the CCC complex, and suggests a model whereby the COMMD-containing proteins were sequentially added to this core complex starting at least as early as the base of tetrapods. If COMMD proteins do indeed function heterogeneously in the complex, this evolutionary scenario would re-capitulate the available structural and biochemical evidence for the complex. In addition, it comports more generally with the increase in the gene content of vesicle trafficking proteins, particulary SNAREs [39, 40, 41] and Rabs [42, 43], at the base of vertebrates.

**Figure 6:**
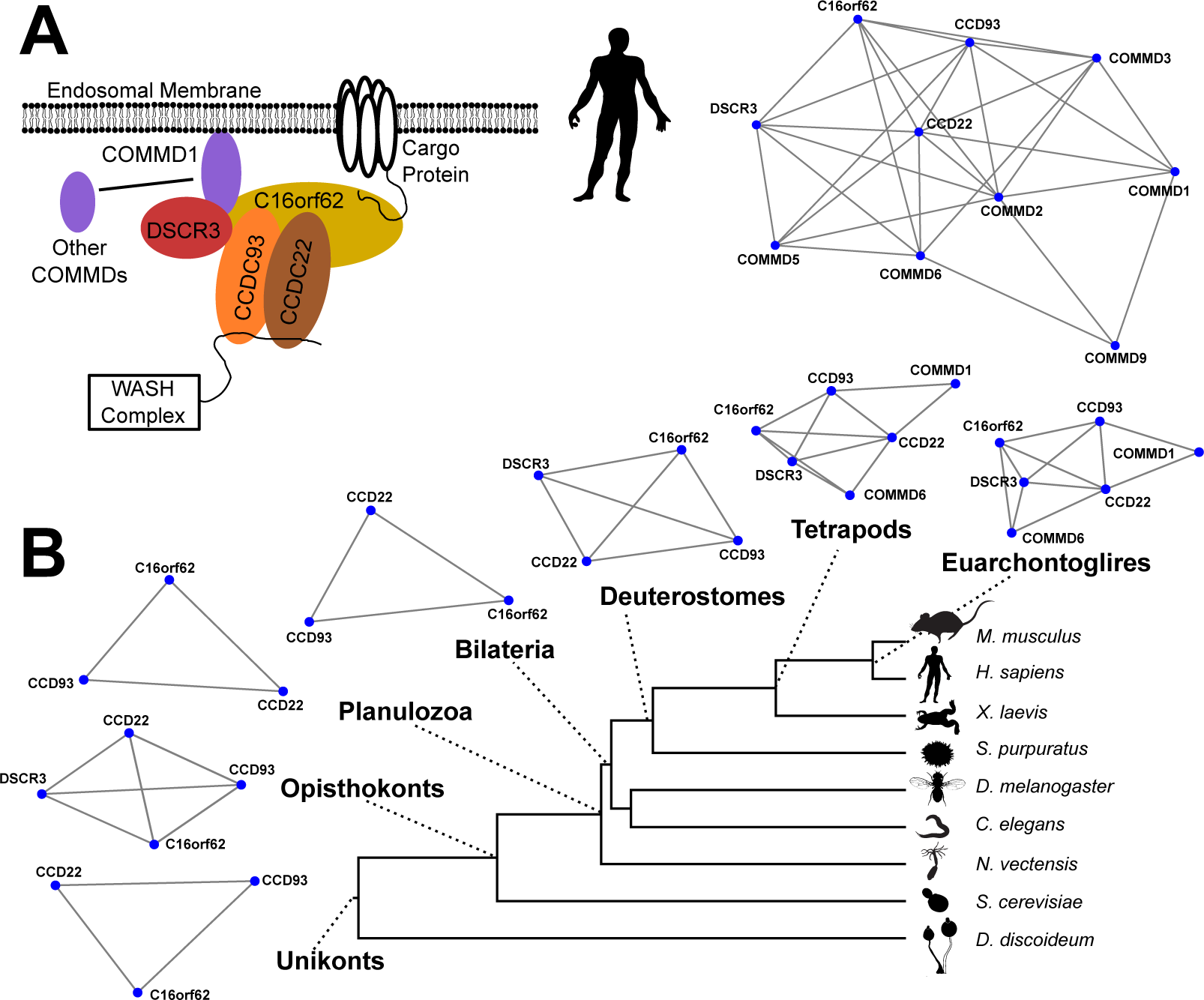
Evolution of the Commander complex. **A.)** Schematic model (Mallam & Marcotte 2017) and PLVM reconstruction of human Commander subunits. **B.)** Interactions between subunits of Commander that survived FDR correction at interior nodes of the tree.

### 2.6. Discussion

We have developed a generalizable phylogenetic latent variable model – a PLVM, which we pronounce “plum” – and have shown that it is capable of performing well on simulated data, of recapitulating known protein complexes across the tree of life, and of reconstructing the evolution of complex protein interactions, all directly from noisy data. Importantly, the model is capable of boosting the power of network inference in species with very poor proteomic sampling by transferring evidence through the phylogeny. We expect this modeling framework and attendant software package to be broadly applicable to any type of comparative network data, and many other types of data besides. With this in mind, we discuss below the important methodological findings and caveats of the model, as well as ways that the model may be extended and improved.

Perhaps the most surprising finding is that fitting the model by likelihood fails to produce adequate predictions, while using an empirical fitting procedure is successful. Our simulations suggest that this is largely driven by the class imbalance problem (Figure 3B), so datasets with balanced classes may perform well under pure likelihood. But when, as in nearly all network applications, true edges are a small fraction of all possible edges, likelihood fails to find the “needles in the haystack.” This “imbalanced learning problem” is well known [44] and plagues most classification procedures. We therefore took the unorthodox step of fitting a generative model using an empirical measure of the recall and precision trade-off called the average precision score (APS). Recall-precision metrics are a standard empirical measure for fitting non-generative classifiers and are preferable in cases of class imbalance [45] but have not, to our knowledge, been used to fit phylogenetic models. This approach improved performance (Figure 3A), but unlike a standard likelihood approach, the output of a model fit under a recall-precision metric cannot be strictly interpreted as a probability. We used a z-score approach that produced adequate results, but model fitting and interpretation will likely be an important area for future improvement.

The results of this model must also be viewed in light of a major artifact that plagues all inferential methods in biology, namely that absence of evidence can appear as evidence of absence. A well-known artifact of comparative methods is that sparse sampling in divergent taxa can lead to the false impression of simplicity both in those taxa and in ancestral nodes [46]. In our case, the tree is highly ladderized and the sampling is sparser on lineages with deeper subtending branches (Figure 1, Table 1), so we cannot rule out that the appearance of “simpler” complexes at deeper nodes, as with the Commander complex, are in fact to due to poor sampling in divergent taxa.

Nevertheless, so little is known about the evolution of protein complexes and molecular systems in general, and so few methods exists for reconstructing the evolutionary process directly from molecular data, that the results are of broad interest. We can conclude, for instance, that the Commander sub-complex comprising the proteins CCDC93, CCDC22, and C16or62 is evolutionarily ancient and was most likely present in the most recent common ancestor of eukaryotes. The data also argue for a complex that included COMMD-containing proteins at least as early as the common ancestor of tetrapods. Notably, this period coincided with an expansion of other protein families involved in vesicular trafficking [39, 42, 43], as well as expansions in receptors and ion channels [47], which is hypothesized to correlate with increased brain complexity as tetrapods invaded dry land [48]. Commander is known to regulate the expression of receptors and ion channels *via* the endosomal pathway [38, 37] and can also affect synaptic function by trafficking copper transporters that regulate metal ion homeostasis in glutamatergic synapses [49, 50]. We can therefore hypothesize that the addition of COMMD-containing proteins to the Commander complex lead to increased cargo-recognition capacity and more complex brain function in early tetrapods. Such hypotheses should be regarded as speculative for the reasons described above, but with ever-expanding datasets the picture will become much clearer.

In addition to increased sampling, one can also envision expansions to the current model. We highlight a few promising areas. First, not all protein-protein interactions evolve at the same rate, and just as with molecular sequence data our data would likely benefit from a modeling framework that allows rate heterogeneity [51]. Secondly, it is likely that using different error models for different taxa would improve model performance. We found that the model performed poorly when yeast co-fractionation data was included. This was likely due to the very poor sampling in yeast (only a single cofractionation experiment is available) relative to other taxa. The model was unable to disregard or down-weight this yeast data because a single error model was used for all taxa. Both of these model expansions may improve performance but will add significantly to parameter complexity. Fitting pro-cedures may also then become a major area of interest, and more theoretical work on these types of models will probably be necessary, especially as regards their identifiability.

In sum, we have demonstrated for the first time that PLVMs are useful for inferring the evolutionary dynamics of protein interaction networks. Many questions in biology, especially those concerning the deep-time evolution of molecular and cellular novelty, are not answerable without a detailed knowledge of the evolutionary processes affecting cellular machines [15]. PLVMs are capable of inferring these processes directly from cutting edge systems data and therefore represent a vital new method for evolutionary systems biology.

## 3. Methods

### 3.1. Data collection and pre-processing

Co-fractionation datasets were derived from Wan *et al.* (2015) and comprise 71 biochemical separations on 9 species (Table 1). Human AP-MS data is from Bioplex2 [52], and yeast from Krogan *et al.* (2006) [34]. In total, our analysis leveraged data from 16774 individual mass spectrometry experiments.

It is not possible to simply compare protein-interaction maps across species because genes can be duplicated and lost. This can be dealt with either by binning proteins into common orthogroups or by explicitly modeling the evolution of interactions across gene duplication and loss events. We chose the former for two reasons. First, gene duplications add substantial complication to the modeling framework and force decisions about how information will be propagated across gene duplication nodes and how low-resolution gene trees will be reconciled with the species tree, both of which are active areas of research in and of themselves. Second, protein-interaction data derives from mass spectrometry, which has lower sensitivity than nucleotide sequencing methods. To determine the abundance of proteins in a sample, proteomics methods match peptide fragmentation spectra against an *in silico* spectrum database derived from a known proteome. Peptide spectra that match multiple proteins are typically removed, so identical or nearly identical regions of closely related paralogs will be ignored. By binning these proteins into or- thogroups, we can use these redundant spectra, thereby boosting sensitivity at the cost of protein specificity.

To convert proteins into orthgroups, we used the eggNOG database and eggNOG-mapper tool [53], which can use both HMMER or DIAMOND for HMM- or sequence-based matching. We matched each protein to its or- thogroup, prioritizing HMMER matches over DIAMOND, and taking only the top hit.

### 3.2. Feature extraction

For each co-fractionation experiment, we extracted two features for every pair of proteins: the Pearson correlation and 1 - the Bray-Curtis distance. The median z-score of these features was then taken across experiments for each species. In addition, we calculated the Pearson correlation between pairs of proteins on a concatenation of all experiments for each species. For AP-MS data, we calculated a hypergeometric p-value for the probability of finding two proteins together in an AP-MS experiment given their abundances across experiments. As a feature, we used the negative log p-value converted to a z-score for each species.

### 3.3. Missing Data

In most phylogenetic methods, missing data comes in the form of gaps in the alignment, and with very few exceptions these gaps are treated as true missing data; that is, as non-informative. But missing data in evolutionary network inference is a subtler problem. There are several reasons why data may be missing and not all should be treated the same way by the model. These are:

1. One orthogroup of the pair is absent in a species.
2. Both orthogroups are absent.
3. Both orthogroups are present but one or both are not observed in the mass spectrometry data.
4. Both orthogroups are observed in some or all of the experiments but one or more features are blank due to pre-processing decisions or other reasons.

Of these, only (3) is completely uninformative missing data. We handle these situations in the following ways. For (1) and (2), pairs of orthogroups where one or both are missing are used as negative training examples during training, and for prediction, the likelihood is set to 0 ensuring that the probability of an interaction at these leaf nodes is set to 0. Situation (3) is treated as missing data, so positive evidence for an interaction in other species is free to propagate to species where evidence is lacking. For situation (4), different features were treated differently. Because the different co-elution features used different filtering thresholds to remove low-abundance proteins, it is possible for one co-elution to be missing while others are present. In this case, we set the missing entry to 0, because missing the abundance threshold is considered evidence of absence. However, the AP-MS features are both sparse and orthogonal to the co-elution features, so if a pair was present in the co-elution data, but not the AP-MS, or *vice versa*, the entries were retained as uninformative missing data. In these cases, the likelihood from multivariate Gaussian is evaluated only on the features that are present.

### 3.4. Training and testing

Training data for the model was taken from two curated protein complex databases: CORUM [31] for human training data and EMBL’s Complex Portal for yeast [32]. These complexes were split into non-overlapping training and test sets, with redundant complexes (those with a Jaccard similarity >.6) being collapsed, ensuring completely independent test and training data. These complexes were then decomposed into all-pairwise positive training examples. Negative training pairs were derived from all pairs of proteins in the positive sets that did not appear together in a complex. For each training set, pairs that included orthogroups known to be missing in other species were annotated as negatives in those species.

We found a good general set of parameters for the simulated annealing algorithm to be:

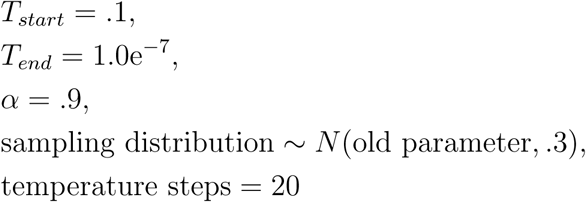

These values for used for fitting for all reported datasets. Each dataset was fit using six independent replicates on one fifth of the training data, and the replicate that performed best on the hold-out four fifths of the training data was chosen for prediction. Prediction was performed on the entirety of the data using the best fit model. The likelihood of interaction between orthogroup pairs where one or both members was missing in a species was set to 0 for that leaf node and all other missing data points was handled as described above.

We then z-scored predictions for all nodes and, using the human old-out data from CORUM, calculated a z-score threshold that corresponded to 25%. This threshold was used for all nodes and only edges above this threshold are reported for Figures 5 & 6. The network shown in Figure 5 was clustered using ClusterONE in Cytoscape with the following parameters: Minimum size: 3; Minimum density: auto; Edge weights: P1 zscore; Node penalty: 2; Haircut threshold: 0; Merging method: multi-pass; Similarity: Match coefficient; Overlap threshold: .8 Seeding method: from unused nodes. The graph layout used was “organic.”

### 3.5. Software and Data Availability

The model is implemented in our Python package “plum” and is available in a Github repository: https://github.com/marcottelab/plum. Data used for the paper is available *via* Zenodo (DOI: 10.5281/zenodo.1406723). We note that the development of this package relied heavily on the publicly available Python modules dendropy [54], pandas [55], scikit-learn [56], and cython [57].

## 4. Extended Methods

### 4.1. Detailed description of the phylogenetic latent variable model

PLVMs are generative models with two components: a component modeling the evolution of the hidden variable, and a component yielding the distribution of observations given the underlying states. In our implementation, the latent variable is a binary state: the presence or absence of an interaction between pairs of orthogroups. We model the evolution of this state as a continuous time Markov chain (CTMC), with each state, i.e. each edge in the network, evolving independently of one another. This framework allows efficient calculation of model fit at the cost of ignoring interactions between edges, and is in keeping with most phylogenetic methods, where each site in an alignment is typically modeled independently. Instead of homologous alignment positions, our model takes in pairs of orthogroups as characters. An edge can take on one of two states: 1, representing an interaction between two proteins, and 0, representing the lack on an interaction. The CTMC model can be fully specified by just two parameters giving the instantaneous rates of change between the two states:

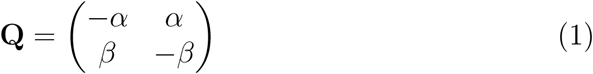

Here, *α* is the rate parameter for 0 *→* 1 transitions (“not interacting” to “interacting”), and *β* is the rate parameter for 1 *→* 0 transitions. The range of the two rate parameters are bounded as 0 *q*_*ij*_ *<*. To determine the probability of observing a change between states over time *τ*, we must derive a probability matrix. For two states, the *P* -matrix can be given in open form [58, 59, 60]:

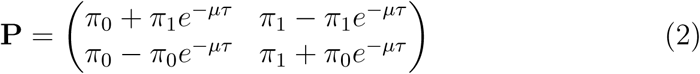

Where *µ* = *α* + *β*. The parameters *π*_0_ and *π*_1_ represent the equilibrium frequencies for states 0 and 1, respectively; that is, the expected frequencies of each state after running the CTMC for an infinite amount of time. These equilibrium frequencies are related to the instantaneous rate parameters as 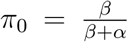 and 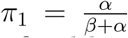 (note that *π*_1_ is just 1 - *π*_0_). Because we con-sider a tree with fixed branch lengths, the rate parameters are the only free parameters for the CTMC component of the model.

This evolutionary framework is exactly the same as those used to describe the evolution of binary morphological characters in standard phylogenetic comparative methods [59, 60]. It suffices for characters whose observations have little or no associated error. However, most functional molecular data, including protein-interaction data, have considerable error associated with them, necessitating an “error” model that translates this uncertainty into a probability of observed data given the latent state. This error model is equivalent to the emission probabilities in a hidden Markov model [29], where each state is associated with its own set of parameters or even its own distribution. Our Python package provides a flexible framework to explore different error models. Single features can be modeled as Gaussian, Gumbel, Cauchy, or Gamma distributions. Due to high false-negative rates in protein-interaction data, we also implement error models where the emission distribution for state 1 is a 2-component mixture, with the lower component capturing false-negatives. However, we envision most applications using several different features, which we model as a multivariate Gaussian, *∼ N* (*µ,* Σ), where covariance matrix Σ can be constrained as a diagonal matrix, or not. Each state, {0, 1}, is associated with its own set of error model parameters that define the expected distribution of continuous data given the presence of that state.

The parameters of the phylogenetic latent variable model to be estimated are then *M* = {*α, β, µ*_0_, Σ_0_, *µ*_1_, Σ_1_}. In addition to these parameters, we denote the topology and branch lengths of phylogeny by *T*, but do not estimate these parameters in our implementation.

### 4.2. Simulation

Because it is a generative model, the full PLVM gives a convenient basis for simulation as well as for probabilistic inference. To simulate a character history, we draw initial states at the root from the equilibrium frequencies, and then traverse the tree from root to tips. State changes along each occur as a Poisson process, where the dwell time in each state is exponentially distributed with parameters drawn from the diagonal entries in the *Q*-matrix. Thus, when the system is in state *i*, the dwell time distribution has rate parameter *-q_ii_* [60]. Dwell times are drawn from the exponential until the additive time exceeds the length of the branch, at which point the state at the node at the end of the branch is set to the current state, and the process beings anew on the child branches. When the process reaches the tips of the tree, the continuous data are drawn from the error model corresponding to the hidden state.

### 4.3. Calculating the tree likelihood

With the error model and the CTMC model in hand, it is now possible to calculate the likelihood of the observed data given a tree and the combined model parameters, *L* = *P* (*D|T, M*). Here, *D* is composed of vectors of one or more features that report on the likelihood of interaction between every pair of orthogroups. Our equivalent to a homologous position in a sequence is an *O × j* matrix for *O* taxa and *j* features. For *N* orthogroups, *D* is then a *N* ^2^ *O× j* matrix; one data matrix for each pair of orthogroups. Because we model the evolution of every edge in the network independently, the total likelihood is the product of the likelihood of each individual edge:

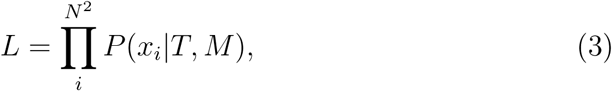

where *x*_*i*_ is the *O × j* feature vector for a pair of orthogroups. To efficiently calculate the likelihood for each *x*_*i*_, we use Felsenstein’s (1981) [30] pruning algorithm, a dynamic programming algorithm that defines a post-order recursion (from the tree tips to root), preventing costly repetitive calculations. Typically, the recursion begins at the leaf nodes by setting the likelihood of observed states to 1 and unobserved states to 0. PLVM models, however, derive the likelihood of each state at the leaves from the error model, *f* (*x*_*i*_). Having been initialized in this fashion, the algorithm goes on to calculate the “conditional likelihoods” at each interior node *n*, which are the likelihoods of all the observations in the clade subtended by the focal node, given that the node is in state *s*. We denote the conditional likelihoods as *L*_*n*_(*s*).

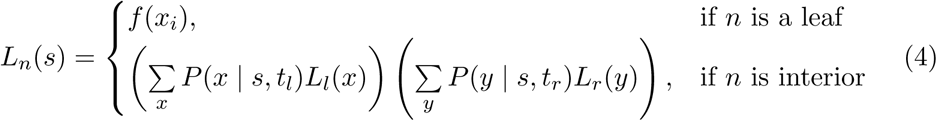

Here, *x* and *y* are the possible states for the left and right child nodes of the focal node *n*, respectively, *L*_*l*_(*x*) and *L*_*r*_(*y*) are the corresponding conditional likelihoods at those child nodes, and the quantities *P* (*x | s, t_l_*) and *P* (*y | s, t_r_*) are the probabilities of observing a change to those states from state *s* over branch length *t*, derived from the *P* -matrix. At the root, the process is terminated and the likelihood of the data is computed by re-weighting the conditional likelihoods for each state by the prior probability of that state, given by the equilibrium frequencies.

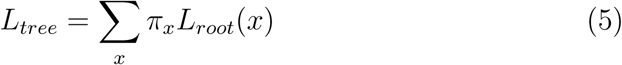

### 4.4. Inferring ancestral states

To infer the probability of each state at each node in the tree, we need two quantities. The first are the conditional likelihoods derived from Felsenstein’s pruning algorithm in the initial post-order trace. The second are found by a pre-order trace (forward, from root to leaves); we will therefore call these the “forward” variables, *F*_*n*_(*s*). The dynamic programming algorithm for these variables is given by Bykova (2013) [29] as

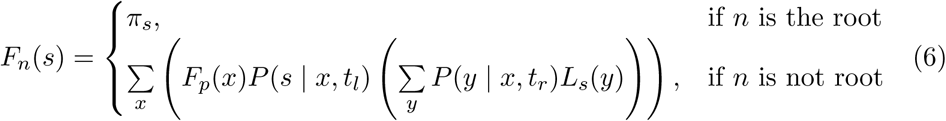

*F*_*p*_ here represents the forward variable in the parent node to *n* and *L*_*s*_ the conditional likelihood at the sister node to *n*. From the forward variables and the conditional likelihoods given by (4), we calculate the probability of ancestral states for each node as:

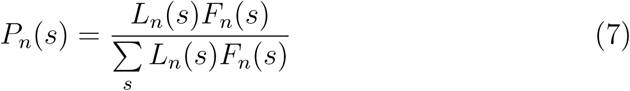

### 4.5. Optimizing the model

We explored two criteria to optimize the model parameters. First, the classical maximum likelihood approach where, given a fixed tree topology and branch lengths, we maximize the probability of the data given the error and network model parameters.

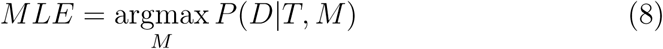

Second, we implemented a supervised learning approach where known interactions are used to derive a classification score for a set of model parameters, again using a fixed tree. Given a set of training protein pairs *D* = [*x*_1_, *x*_2_*…x_N_*] where the state ∈ {0, 1} is known from curated databases, we infer ancestral probabilities of an interaction and then rank the pairs by their probability of interaction. From this ranking, the recall, *R*, and precision, *P*, can be calculated at each threshold *t* from the number of true positives *TP* and false positives *FP*, as 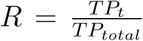 and 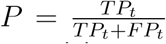 We then use the av-erage precision score of the whole recall-precision curve as our optimization criterion:

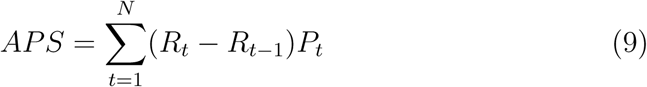

This calculation is implemented using the scikit-learn python package [56]. For both the likelihood and the *APS* scores, we found optimal sets of model parameters using a simulated annealing algorithm. The algorithm begins at a certain “temperature,” *T*, and steadily decrements it by a fraction *α* where 0 *≤ α <* 1. At each temperature, a new parameter value is sampled stochastically from a Guassian distribution centered on the previous value and the optimization score is calculated. If the new score is higher than the previous one, the new parameter value is kept, otherwise, the new value is picked with probability 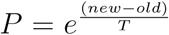 In our Python package, the user can vary the starting temperature, the ending temperature, the decrementing factor, the standard deviation of the Gaussian sampling distribution, and the number of sampling steps at each temperature.

## 5. Acknowledgements

The authors gratefully acknowledge Kevin Drew, Anna Mallam, and Claire McWhite for assistance and critical suggestions, Ammon Thompson and Michael Landis for helpful discussions about the modeling framework, the Texas Advanced Computing Center at the University of Texas for high-performance computing resources, and funding from the National Institutes of Health (5F32GM112504-03 to B.J.L and R01 HD085901, R35 GM122480, R01 DK110520 to E.M.M.), National Science Foundation (IOS-1237975, to E.M.M.), and Welch Foundation (F-1515, to E.M.M.).

